# The selectivity implications of docking libraries with greater and lesser similarities to bio-like molecules

**DOI:** 10.64898/2026.02.02.703317

**Authors:** Brendan W Hall, Kensuke Sakamoto, Xi-Ping Huang, John J Irwin, Brian K Shoichet, Bryan L Roth

## Abstract

As virtual libraries have expanded into the tens of billions via make-on-demand chemistry, their similarity to metabolites, natural products, and drugs (“bio-like” molecules) has rapidly diminished. Despite this divergence, molecular docking of these ultra-large libraries has yielded molecules at higher experimental hit-rates and with improved affinities. The structural divergence from bio-like space raises the possibility that molecules from these ultra-large libraries have improved selectivity. Just as plausibly, if hit-rates on-target are divorced from similarity to bio-like molecules, so too may be selectivity against off-targets. Here, we test whether docking hits for the 5-HT_2A_ serotonin receptor from ultra-large libraries are more selective than those from smaller and more bio-like “in-stock” libraries. Chemoinformatic similarity predicts that docking actives from the in-stock library have more off-targets than the more chemically novel hits emerging from docking the ultra-large library. This may reflect the bias of the known, however, as when tested experimentally at scale against 318 GPCRs, both 16 agonists from the ultra-large library and 20 actives from the in-stock library had similar numbers of off-targets. While the ultra-large library hits are more sub-type selective for the 5-HT_2A_ over the 5-HT_2B_ and 5-HT_2C_ receptors, overall these results may suggest that selectivity against off-targets, like affinity and hit-rates for on-targets, is divorced from library similarity to bio-like molecules.

## Introduction

Make-on-demand (“tangible”) libraries^1, 2^ have expanded readily-available chemical space to nearly 80 billion molecules (Enamine REAL accessed Jan 2026). Docking subsets of these ultra-large libraries has led to potent and selective chemical probes for diverse protein classes, including GPCRs^3–9^, enzymes^10–14^, transporters^15^, and others^16, 17^. Because these libraries are constructed combinatorially from diverse synthetic building blocks, their growth has led to decreasing similarity to “bio-like” molecules (metabolites, natural products and drugs)^18^. Whereas similarity to bio-like molecules has been thought to be important for the success of compound screening^19–22^, both computational and experimental, instead docking of these less-and-less bio-like libraries yields better-fitting and better-scoring molecules than smaller, more bio-like libraries^3, 18, 23^. Tested experimentally at scale, docking hits from the new libraries have higher hit-rates and affinities than those from smaller more bio-like libraries^3, 23^.

One might expect docking hits from ultra-large, less bio-like libraries to be more selective than those from smaller, more bio-like libraries. After all, many biologically relevant ligands, such as neurotransmitters, interact with multiple receptors. Serotonin (5-hydroxytryptamine, 5-HT) engages 14 receptors across seven classes^24, 25^, while dopamine acts through five distinct receptors^26^. Some receptors are even more promiscuous; for instance, trace amine-associated receptor 1 (TAAR1) can be activated by trace amines like tryptamine and octopamine, as well as by dopamine and to a lesser extent serotonin^27–29^. Promiscuity across bio-like molecules also extends across receptor families: dopamine can activate adrenergic receptors^30, 31^ and norepinephrine has higher potency for D4- dopamine receptors than does dopamine itself^32, 33^. Molecules from ultra-large libraries that diverge from bio-like chemical space could be less prone to this promiscuity and exhibit more selective binding. Conversely, if promiscuity, like affinity and hit rate, is unrelated to the similarity of the library to bio-like molecules, then actives from docking the ultra-large libraries may be no more selective than those from the in-stock libraries. Distinguishing between these hypotheses demands systematic comparison of selectivity between ultra-large and smaller libraries.

Here, we computationally and experimentally compared the selectivity of docking hits for 5-HT_2A_ serotonin receptor agonists from two chemical libraries differing in bio-likeness. One library was a 1.6 billion molecule make-on-demand library that was far less bio-like than a second 3.5 million molecule in-stock library. We docked the in-stock library against an agonist bound 5-HT_2A_ structure and collected ultra-large library docking results from a previous study^4^. We computationally assessed off-target binding with the Similarity Ensemble Approach (SEA)^34, 35^ and experimentally tested docking hits from each library for 5-HT_2A_ binding and functional activity across all 5-HT_2_ -family receptors. We also profiled key compounds for agonist activity at the druggable GPCR-ome using the PRESTO-Tango resource^36^, and performed radioligand binding studies to quantify binding affinities against a subset of aminergic GPCRs and related proteins.

## Results

### Bio-likeness of the ZINC and in-stock libraries

We first quantified the diversity and bio-likeness of two virtual screening libraries: a 1.6 billion molecule make-on-demand library used by Lyu et al^4^, subsequently referred to as the “ZINC library”, and a smaller 3.5 million molecule informer set of in-stock molecules from ZINC20^37^, subsequently referred to as in-stock. Following prior work^18^, we calculated the maximum similarity of molecules from each library to a reference set of bio-like molecules (metabolites, natural products, and drugs). The in-stock library was more bio-like, with an average Tanimoto Coefficient (Tc) of 0.37 (ECFP4 fingerprints) compared to 0.31 for the ZINC library (**Fig. 1A**). To put this in perspective, random similarity for this fingerprint occurs at around Tc of 0.25, and a Tc of less than 0.35 is considered a scaffold-hop^38^. Perhaps more important than the average Tc difference to bio-like molecules, there was a large skew in the in-stock library toward bio-like molecules (Pearson moment coefficient of skew of 2.33)—visible in the long tail in **Fig. 1A**—which was far less apparent in the ZINC library (skew of 1.1). More intuitively, 54% of the in-stock library molecules had Tcs >0.35 to at least one bio-like molecule, while only 17.2% of the ZINC molecules did, and 6.3% of the in-stock molecules had Tcs of >0.5—a value indicating high similarity—to bio-like molecules, 20-fold more than the percentage of ZINC molecules that did so.

**Figure 1.**
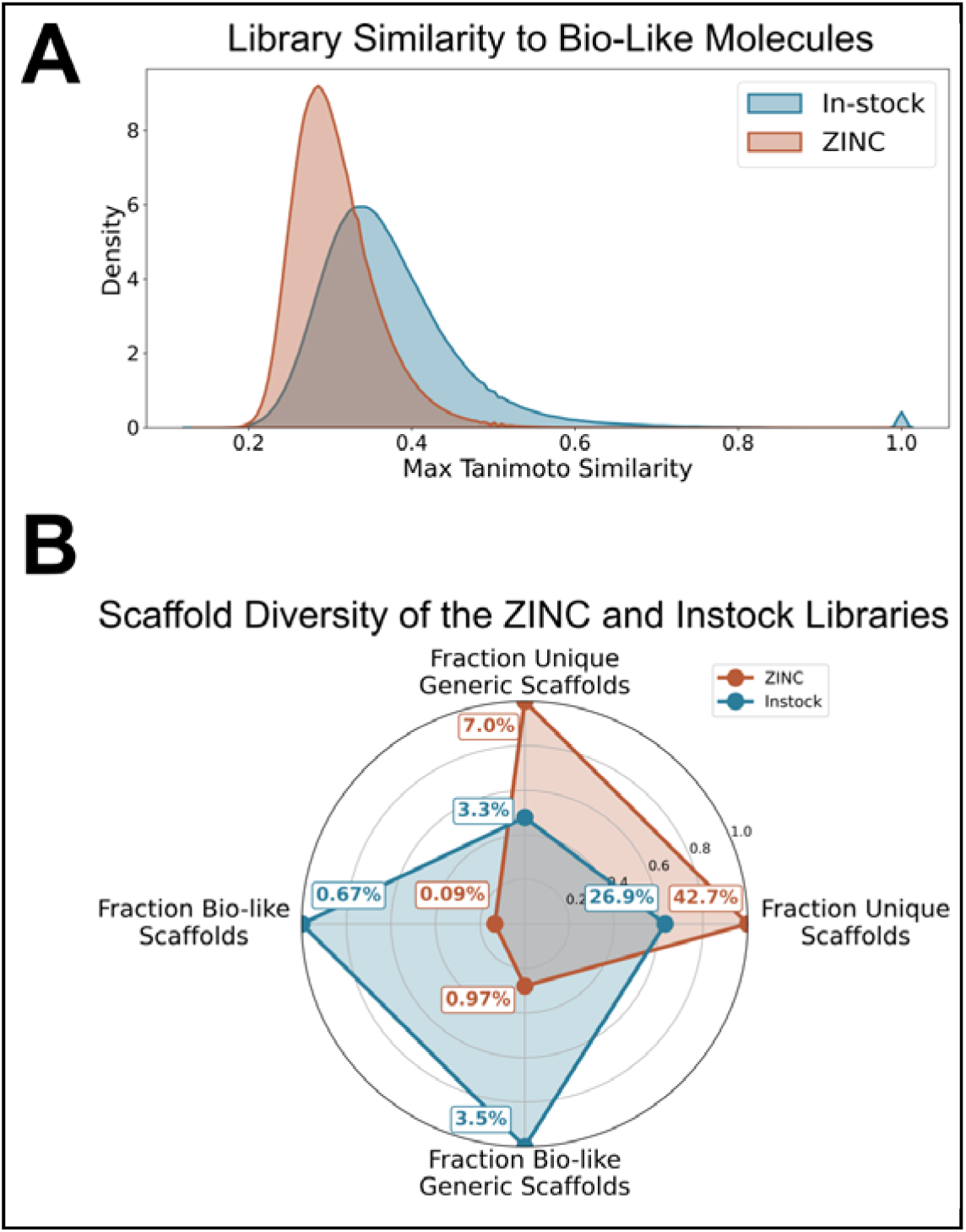
Library bio-likeness and diversity analysis. **(A)** Distributions of maximum Tanimoto similarity (ECFP4 fingerprints) between in-stock and ZINC molecules and a set of bio-like molecules. **(B)** Fractions of unique standard and generic Bemis-Murcko scaffolds in the in-stock and ZINC libraries, and the fractions of each scaffold type shared with the bio-like set.

The in-stock library was also less diverse and shared more scaffolds with the bio-like set (**Fig. 1B**). Only 26.9% of in-stock molecules contained a unique scaffold, compared to 42.7% in ZINC. Strikingly, in-stock scaffolds were 7.4-fold more likely than ZINC scaffolds to overlap with bio-like scaffolds. If selectivity among receptors was driven by similarity to bio-like molecules, which typically bind at least two targets and for metabolites and transmitters often many more^39, 40^, then we might expect docking hits from in-stock libraries to be more promiscuous than those from ultra-large libraries like ZINC. Conversely, if promiscuity and selectivity were more a function of receptor recognition and how it related to that of other targets, then the selectivity of docking hits from the two libraries might be more similar. This we set out to test for docking hits from both libraries against the 5-HT_2A_ receptor, measuring selectivity for this target against 318 other receptors in the GPCR-ome.

### Workflow for comparing the selectivity of ZINC and in-stock docking hits

We tested whether ZINC docking hits are more selective than those from the in-stock library by both computation and experiment (**Fig. 2**). We used ZINC 5-HT_2A_ docking results from a previous study^4^ and conducted a new docking screen of the in-stock library, using the same sampling, scoring, and prioritization (see Methods). After inspecting 2,526 in-stock molecules that passed filtering, we ordered 85 for testing; we compared these to the 384 ZINC molecules from a previous study^4^.

**Figure 2.**
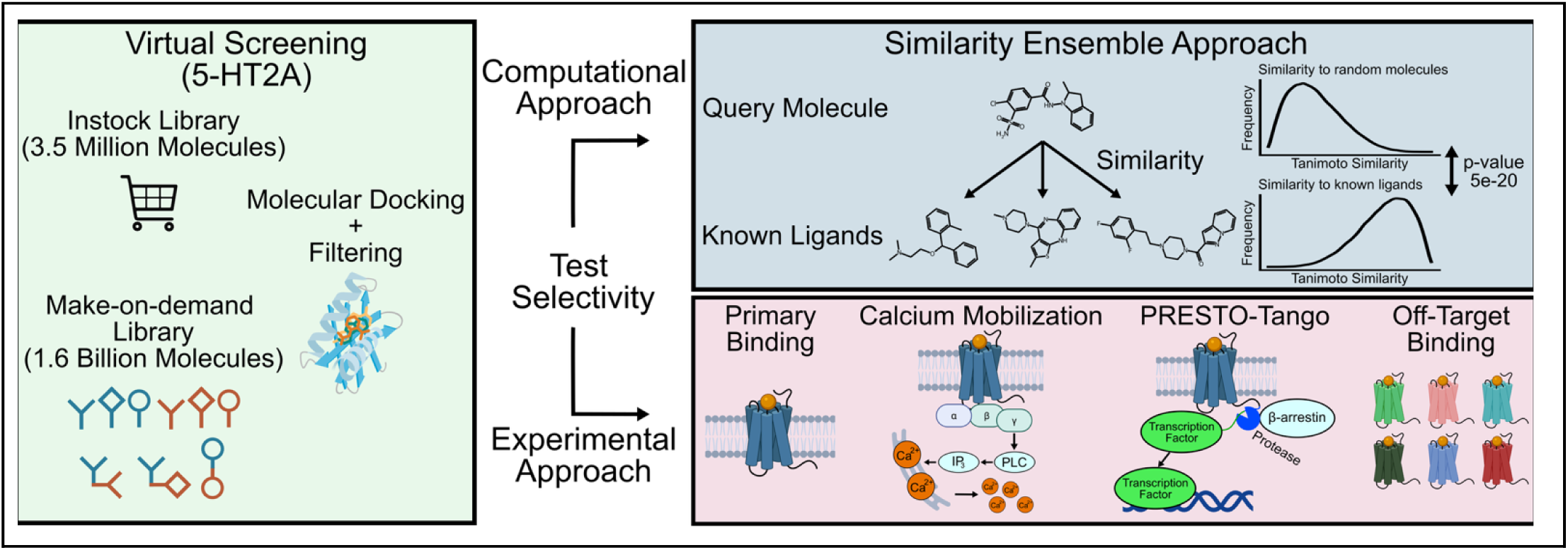
Computational and experimental workflow to test the selectivity of the in-stock and ZINC libraries. We docked both libraries against 5-HT_2A_ and selected top-ranking molecules for computational and experimental selectivity testing. Computationally, we used the Similarity Ensemble Approach (SEA) to predict protein targets. Experimentally, we measured 5-HT_2A_ binding with radioligand displacement, 5-HT_2A/B/C_ agonism and antagonism with calcium mobilization, agonist activity across the druggable GPCR-ome with PRESTO-Tango, and off-target GPCR binding with radioligand displacement.

Computationally, we used the Similarity Ensemble Approach (SEA)^34, 35^ to predict protein off-targets for the top 10,000 scoring molecules and for the synthesized docking hits from each library. Experimentally, we evaluated 5-HT_2A_ binding using radioligand displacement assays, performing new experiments for the in-stock docking hits and using results from a previous study^4^ for the ZINC hits. We quantified functional activity with calcium mobilization assays across all 5-HT_2_-family receptor subtypes and collated prior data for ZINC hits. To investigate the broader promiscuity for both libraries, we profiled a subset of molecules from each library across 318 GPCRs using the PRESTO-Tango assay^36^. Finally, we determined off-target binding affinities against a set of 21 aminergic GPCRs and related proteins^41^. Experimental details are provided in the Methods.

### SEA off-target predictions for the ZINC and in-stock docking hits

We used the Similarity Ensemble Approach (SEA)^34, 35^ to computationally predict protein off-targets of docking hits from the ultra-large ZINC library and the smaller in-stock library. SEA predicts potential protein targets for a query molecule by comparing its chemical similarity to sets of known ligands from ChEMBL^42^ and determining whether these similarities are greater than expected by chance, given a random background. We generated SEA predictions for the top 10,000 molecules ranked by docking score (**Data. S1)** from each library, and the molecules prioritized for experimental testing (**Data. S2)**. We considered SEA predictions with p-values less than (better than) 10^-10^ as substantial. Because our docking campaign targeted an aminergic GPCR—the 5-HT_2A_ receptor—we focused on SEA predictions for aminergic GPRCs and related proteins (**Table. S1**).

Of the top 10,000 scoring docked molecules, SEA predicted 4,316 from the in-stock library to bind at least one aminergic GPCR, compared to 652 from the ultra-large ZINC library (**Fig. 3A and Data. S3**). SEA also predicted in-stock molecules to bind more aminergic GPCR off-targets than it did for ZINC molecules (**Fig. 3B**). For molecules with at least one off-target, in-stock molecules had an average of 4.8 predicted off-targets versus 2.1 predicted for the ZINC molecules. We observed similar trends among the molecules prioritized for testing. SEA predicted 49.4% of in-stock hits to bind an aminergic GPCR versus 25.5% predicted for the ZINC hits (**Fig. 3C and Data. S4**), with 3.9 average predicted off-targets for the in-stock hits versus 2.2 for the ZINC hits (**Fig. 3D**). These differences persisted across more stringent SEA p-value thresholds (**Fig. S1 and Fig. S2**).

**Figure 3.**
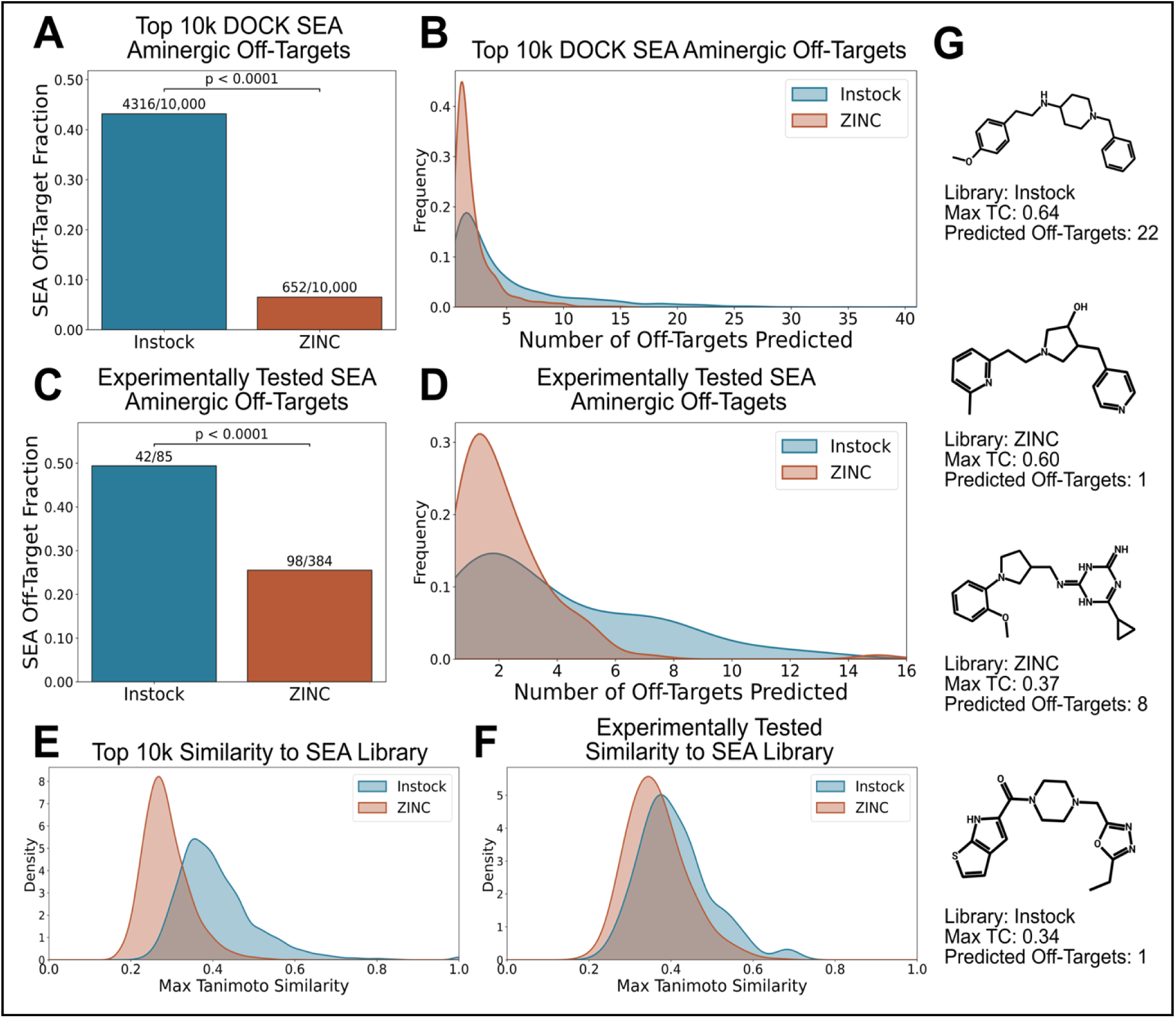
Similarity Ensemble Approach (SEA) predictions for in-stock and ZINC molecules. **(A)** Fraction of the top 10,000 DOCK-scoring molecules predicted by SEA to bind any aminergic GPCR. **(B)** Number of SEA off-target predictions for the top 10,000 molecules. **(C)** Fraction of experimentally tested molecules predicted to bind any aminergic GPCR. **(D)** Number of SEA off-target predictions for the experimentally tested molecules. **(E)** Similarity of the top 10,000 molecules to the SEA library of known aminergic GPCR binders. **(F)** Similarity of the experimentally tested molecules to the SEA library of known aminergic GPCR binders. **(G)** Representative molecules from each library with their similarity to the SEA library and number of predicted off-targets.

While SEA predicted a large difference in selectivity between the two libraries, we anticipated that the predictions simply reflected similarities to known aminergic GPCR ligands in ChEMBL. Indeed, in-stock molecules were more similar to the set of active ligands for aminergic GPCRs with an average maximum ECFP4 T_C_ of 0.41, versus the 0.29 Tc for ZINC molecules (**Fig. 3E and Fig. 3F**), a substantial difference for this fingerprint^38^. However, similarity alone did not account for promiscuity, evidenced by high similarity compounds with few predicted off-targets and low similarity molecules with many predicted off-targets (**Fig. 3G**). Furthermore, the greater predicted promiscuity of in-stock molecules extended beyond aminergic GPCRs to other SEA targets (**Fig. S3**). Because the SEA predictions were biased toward known chemical space, we turned to experimentation to further test the selectivity of the two libraries.

### 5-HT2A primary binding and 5-HT_2_ receptor family functional activity of ZINC and in-stock docking hits

We compared the 5-HT_2A_ binding of top-scoring molecules from the in-stock and ZINC libraries via radioligand binding assays. Intriguingly, hit-rates were similar between the libraries. For the in-stock molecules, 20 of 85 displaced more than 50% of [^3^H]-LSD at 10μM (23.5% hit rate; **Fig. 4B**; **Fig. 4C and Data. S5**), while 93 of 384 ZINC molecules did so in a previous study (24.2%; **Fig. 4A and Fig. 4C**)^4^.

**Figure 4.**
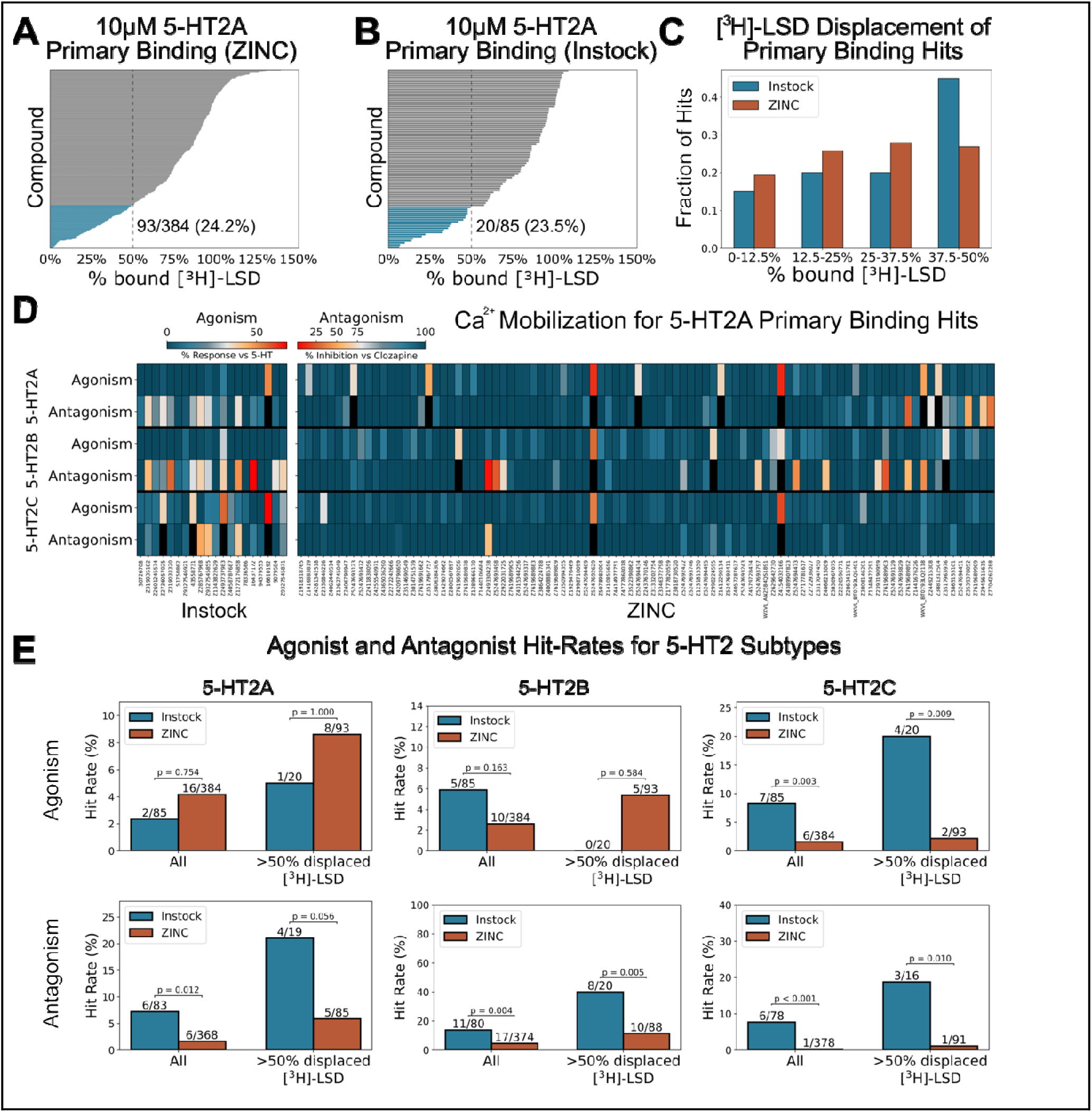
5-HT_2A_ Primary binding screening and 5-HT_2A/2B/2C_ calcium mobilization assays. Fraction of [^3^H]-LSD displaced by **(A)** ZINC and **(B)** in-stock molecules at 10μM. The dashed line indicates 50% displacement and the values show the number of molecules with >50% displacement. **(C)** Binned fractio s of [^3^H]-LSD displacement by 10μM of molecules from both libraries showing those with >50% displacement. **(D)** Calcium mobilization assay in agonist and antagonist modes, with compounds tested at 3μM. Heatmap coloring is scaled such that 10% of 5-HT agonist activity and 20% of clozapine antagonist activity corresponds to the midpoints. Only molecules with >50% [^3^H]-LSD displacement are shown. **(E)** Agonist and antagonist hit rates for in-stock and ZINC molecules vs. 5-HT_2A_, 5-HT_2B_, 5-HT_2C_. Agonist hits were defined as >10% of 5-HT activity; antagonist hits were defined as >20% of clozapine activity. ZINC data for all plots comes from previous work^4^.

We next assessed the functional activity of molecules from each library across the 5-HT_2_ receptor family using a calcium mobilization assay. We tested molecules at 3 μM for both agonism, defined as ≥10% 5-HT response, and antagonism, defined as ≥20% clozapine inhibition. Despite docking to an agonist-bound 5-HT_2A_ structure, we found a mix of agonists and antagonists, something we have observed previously with many GPCRs^1, 4, 6, 43, 44^; here in-stock molecules were actually more likely to be 5-HT_2A_ antagonists than agonists, with a ratio of agonists to antagonists of 2:6 (**Fig. 4D**; **Fig. 4E and Data. S6**). The ZINC molecules had higher fidelity to the active state of the receptor, with the ratio of agonists to antagonists discovered rising to 16:6.

This higher functional selectivity for the ZINC molecules against 5HT_2A_ was paralleled by subtype selectivity within the 5HT_2_ family. In-stock molecules were more likely than ZINC molecules to show activity at 5-HT_2B_ and 5-HT_2C_ receptors, which are frequent off-targets of 5HT_2A_ -preferring molecules. 18.8% of in-stock molecules showed functional activity at the 5-HT_2B_ receptor, compared to 7.0% for ZINC. For 5-HT_2C_, 15.3% of in-stock molecules showed activity, versus only 1.8% for ZINC. Strikingly, these differences were more pronounced among the 5-HT_2A_ primary binding screening hits, although interestingly none of the primary binding in-stock hits were 5-HT_2B_ agonists (**Fig. 4E**). These trends remained with stricter definitions of agonism and antagonism (**Fig. S4**).

### PRESTO-Tango screen of ZINC and in-stock docking hits

Because in-stock molecules were more functionally promiscuous at 5-HT_2_ -family receptors, we next asked whether they were also more likely to have activity at other GPCRs. To address this question, we used the PRESTO-Tango resource^36^ which employs β-arrestin recruitment assays across 318 GPCRs representing most of the druggable GPCR-ome. Here we profiled 20 in-stock molecules, which were selected for >50% [^3^H]-LSD displacement and 16 ZINC molecules which were selected as 5-HT_2A_ agonists. We classified molecules as active if they produced greater than a 3-fold increase in activity over basal levels.

Despite increased promiscuity within the 5-HT_2_ family, in-stock molecules did not differ from ZINC molecules in activities against 318 GPCRs. Both libraries had broad off-target activities, with in-stock molecules active at 0.3%-4.4% of receptors (mean 1.7%) and ZINC molecules at 0-6.6% (mean 2.1%) (**Fig. 5A**; **Fig. 5C, and Data. S7**). Because the docking screens targeted an aminergic GPCR, we next investigated activity across a subset of 55 aminergic GPCRs. Here too, we observed no significant differences between molecules from the two libraries and both libraries showed wider ranges of off-target activities, with in-stock molecules active at 0%-11% of aminergic receptors versus 0%-25.5% for ZINC molecules (**Fig. 5B and Fig. 5D**). Indeed, the most promiscuous compounds in this GPCR profiling assay were found in the ZINC library. The lack of difference between libraries persisted with a stricter definition of off-target activity (**Fig. S5**).

**Figure 5.**
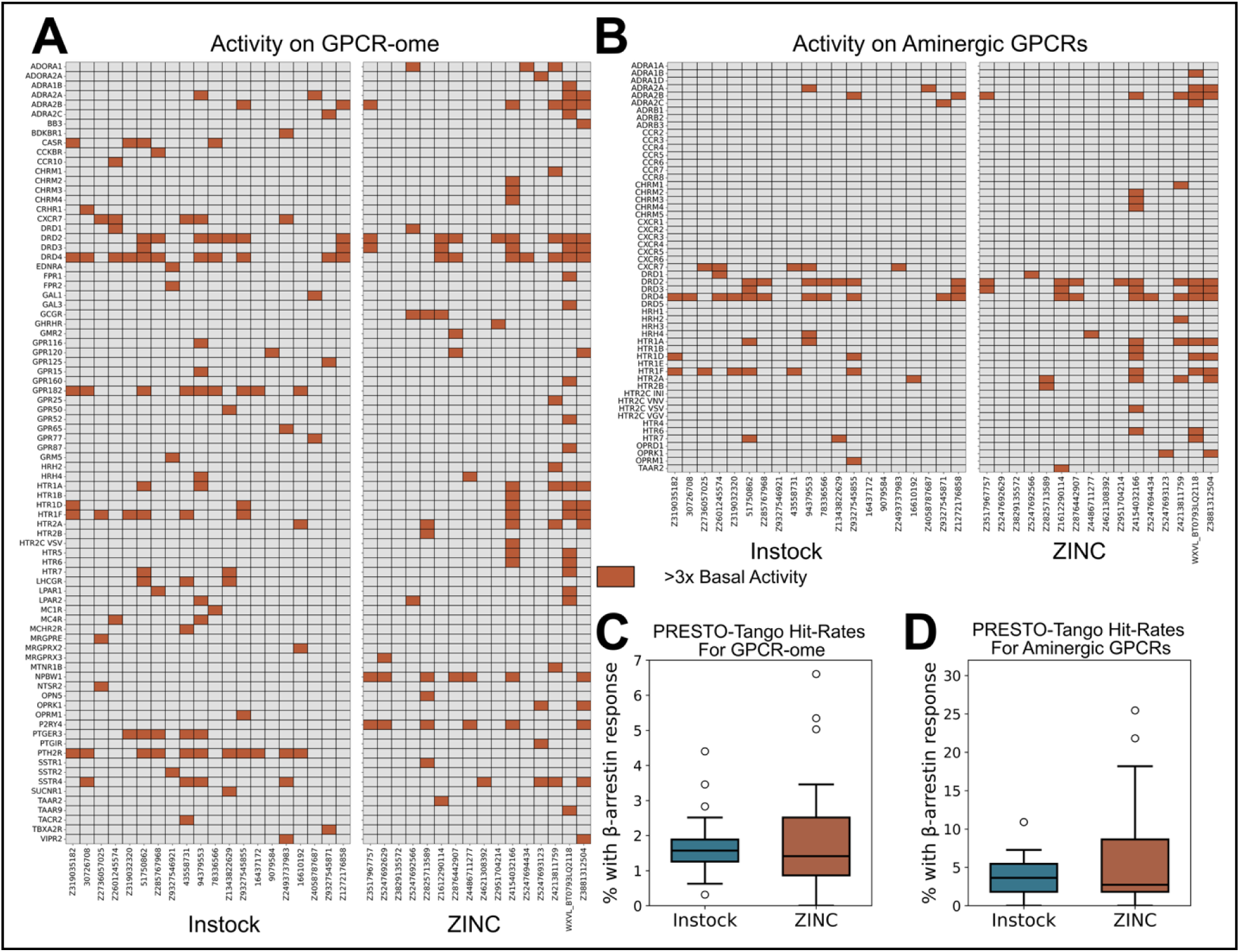
Druggable GPCR-ome activity. PRESTO-Tango hits for in-stock and ZINC molecules across the **(A)** druggable GPCR-ome and the **(B)** aminergic GPCRs. Molecules were considered hits if they produced a >3-fold increase in activity over basal levels. PRESTO-Tango hit rates for in-stock and ZINC molecules across the **(C)** druggable GPCR-ome and the **(D)** aminergic GPCRs.

### Aminergic GPCR binding affinity of ZINC and in-stock docking hits

We next asked whether the two libraries differed in off-target binding to a subset of aminergic GPCRs and related proteins. Here we measured off-target binding by radioligand displacement at 10μM of compound (**Fig. 6A and Data. S8**). There was no significant difference in off-target hit rates (defined as >50% displacement) between the libraries, but both libraries showed broad distributions (**Fig. 6B**). In-stock molecules had hit rates ranging from 4.8% to 61.9% (average of 30.1%), while ZINC molecules ranged from 4.8% to 90.5% (average of 28.5%). Interestingly, the most promiscuous compound came from the ZINC library despite having a relatively low bio-likeness (Tcmax=0.29). Although molecules from both libraries tested in this assay showed similar functional hit-rates at 5-HT_2B_ and 5-HT_2C_ (40% and 35% for in-stock; 44% and 38% for ZINC), they differed in primary binding. In-stock molecules had binding hit rates of 85% and 10% for 5-HT_2B_ and 5-HT_2C_, respectively, compared to 69% and 25% for ZINC (**Fig. 6C**).

**Figure 6.**
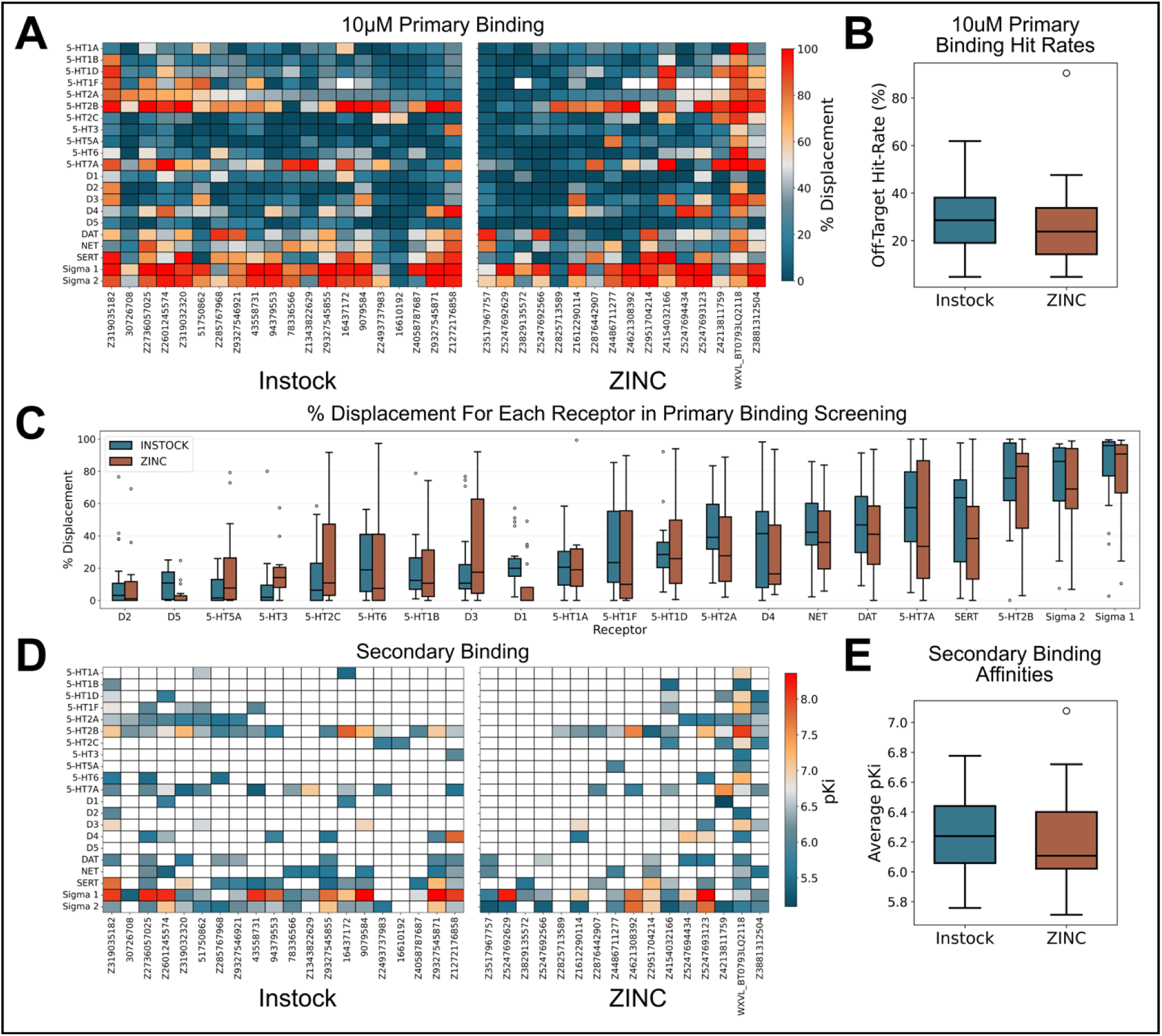
Off-target aminergic GPCR binding. **(A)** Fraction of radioligand displaced by 10μM of in-stock and ZINC molecules across a subset of aminergic GPCRs and related proteins. **(B)** Hit rates for in-stock and ZINC molecules against this subset. Molecules were considered hits if they displaced >50% of the radioligand. **(C)** Radioligand displacement by 10μM in-stock and ZINC molecules for each aminergic GPCR tested. **(D)** pKi values for in-stock and ZINC molecules that displaced >50% of radioligand in aminergic GPCR primary binding assays. **(E)** Average pKi values of in-stock and ZINC molecules in secondary binding assays.

Lastly, we asked whether the binding affinities of the hits differed between libraries. For each compound, we determined pK_i_ values for aminergic GPCRs where radioligand displacement exceeded 50% in the primary binding assay (**Fig. 6D and Data. S9**). Average off-target pK_i_ values were similar between in-stock and ZINC molecules with the in-stock molecules having an average pKi of 6.3 versus 6.2 for the ZINC molecules (**Fig. 6D and Fig. 6E**).

## Discussion

The growth of make-on-demand chemical libraries has been accompanied by decreasing similarity to bio-like molecules (natural products, metabolites, and drugs). Counter-intuitively, docking hits from these expanding libraries show improved docking scores, hit rates, and affinities compared to those from smaller, more bio-like libraries, belying previous inferences that high bio-likeness is important for the success of library screening methods^19–22^. Topologically unrelated molecules can bind receptors using physically similar groups, and as the libraries grow more compounds that physically complement a given target are sampled, partly explaining the higher hit-rates and affinities from the larger libraries. A plausible extension of this observation is that docking ultra-large libraries will also lead to more selective molecules, not just more potent ones. Just as plausible, however, is that if similarity to bio-like molecules does not correlate with hit-rates and affinities from virtual screens, it might not correlate with selectivity either.

Looking at selectivity for the 5-HT_2A_ receptor of actives from a large make-on-demand library (ZINC) versus a much smaller in-stock library, our results more strongly support the second hypothesis. Against 318 other GPCRs, there was little selectivity difference between docking experimental actives from the in-stock and ultra-large libraries. Admittedly, we did observe more functional and sub-type selectivity for the ultra-large library actives, which were more selective as 5-HT_2A_ agonists than those from the in-stock library. The in-stock actives were more likely than the ultra-large library actives to be 5-HT_2A_ antagonists and to be active at the 5-HT_2B_ or 5-HT_2C_ receptor off-targets. While this sub-type and functional selectivity is encouraging, and may be a harbinger for other targets, it must be admitted that they represent a carve out of our overall results, and are supported by smaller numbers, if still statistically significant ones. For instance, while the ratio of agonists to antagonists was meaningfully higher for the ultra-large library screen than for the in-stock screen, the percent of agonists found among the molecules tested from both screens did not statistically differ. Broadly, the results of this study more strongly support the idea that target selectivity is not a function of similarity to bio-like molecules within libraries.

Some cautions deserve emphasis. Because we tested fewer in-stock docking hits, and these were less likely to be 5-HT_2A_ agonists than ZINC docking hits, we used slightly different selection criteria for the PRESTO-Tango and secondary binding assays. For the ZINC library, we selected molecules identified as 5-HT_2A_ agonists in the calcium mobilization assay. For the in-stock library, this did not yield enough molecules, so we instead selected molecules that displaced >50% [^3^H]-LSD in 5-HT_2A_ primary binding screening. Furthermore, ZINC docking hits came from our previous work, where we performed two docking screens. One used a cryo-electron microscopy structure of 5-HT_2A_ bound to lisuride (the structure used for the in-stock docking screen in this study), and the other used an AlphaFold2^45^ predicted structure. Differences in criteria for selection for secondary screening or docking setup may have influenced the observed off-target binding. Finally, this study, while relatively extensive in docking and even in number of compounds tested, represents results from only one target—more research is warranted in this area.

These caveats should not diminish the main findings of this study. Docking hits against the 5-HT_2A_ receptor from ultra-large virtual libraries, despite their increasing divergence from bio-like chemical space, exhibited distinct selectivity profiles. Compared to hits from the smaller, more bio-like library, hits from the ultra-large library were more selective as 5-HT_2A_ agonists, showing reduced activity at 5-HT_2B_ and 5-HT_2C_. In contrast, both libraries produced hits with comparable selectivity across the broader druggable GPCR-ome and a subset of aminergic GPRCs. Whether further growth of make-on-demand libraries and docking campaigns by additional orders of magnitude will yield molecules with improved selectivity across the GPCR-ome remains an open and important question.

## Methods

### Bio-likeness + Scaffolds Calculation

Following Lyu, Irwin, and Shoichet^18^, we used the ZINC15^46^ worldwide drug set (5,900 molecules) and the ZINC15 biogenic set (168,185 molecules) to represent bio-like chemical space. We quantified the bio-likeness of a molecule by its maximum Tanimoto coefficient to any molecule in this bio-like set. We used Morgan fingerprints generated with rdkit with a radius of 2 and a bit length of 2,048. We used rdkit to generate Bemis-Murcko scaffolds and generic Bemis-Murcko scaffolds^47^ for the ZINC and in-stock libraries. We considered scaffolds bio-like if they appeared in the set of bio-like molecules. We considered all 3,490,488 in-stock molecules and an equivalently sized, representative set of the ZINC22 ultra-large library.

### Docking Against 5-HT_2A_

We collected 1,620,934,121 ZINC22^48^ ultra-large library docking results against a cryo-electron microscopy 5-HT_2A_ structure from a previous publication^49^, and applied a similar filtering workflow here. Using the same docking setup and DOCK3.8, we docked 3,490,488 molecules from ZINC22g, a subset of the ZINC22 library containing in-stock molecules for rapid acquisition, against 5-HT_2A_. On average, docking explored 258 orientations and 194 conformations per molecule. We collected 29,771 top-ranking molecules with docking scores better than −40.32, the score threshold used in our previous study. Following that workflow, we filtered these molecules for novelty against 8,601 known 5-HT_2A_, 5-HT_2B_, and 5-HT_2C_ ligands, resulting in 24,664 molecules. We then applied two interaction filtering procedures. The first required interactions with D155, S242, S239, T160, S159, and F340, exactly as in our previous study, resulting in 105 molecules. The second required only interactions with D155 and F340, resulting in 3,816 molecules. We clustered the remaining molecules using LUNA^4^ 1,024 bit fingerprints with a Tanimoto coefficient threshold of 0.35, yielding 73 and 2,526 cluster heads, respectively. From the first filtering method, we ordered 37/73 of the cluster heads and received all 37. From the second filtering method, we visually inspected the docking poses of the 2,526 cluster heads for overall shape complementarity and ordered 50, of which we received 48.

### Similarity Ensemble Approach (SEA) Calculations

The Similarity Ensemble Approach (SEA)^34, 50^ predicts protein binders for a query molecule by evaluating whether its similarity to known ligands differs significantly from its similarity to random molecules. We used tldr.docking.org to make SEA predictions using CHEMBL21^35^ for the top 10,000 DOCK-scoring molecules from the in-stock and ZINC docking screens, as well as for the experimentally tested molecules. We considered SEA predictions with p-values below 1e-10 as significant and retained only aminergic GPCRs (Table. S1).

### 5-HT2A radioligand binding

Competition binding assay was performed with human 5-HT_2A_ receptors transiently expressed in HEK293 cells and prepared as crude membrane fractions in a 96-well format. Assay conditions were adapted from NIMH Psychoactive Drug Screening Program (PDSP) protocols (https://pdsp.unc.edu/pdspweb/?site=assays). Test compounds were supplied as 10 or 100 mM stocks in dimethyl sulfoxide (DMSO) and diluted into standard binding buffer [50 mM Tris-HCl (pH 7.4), 10 mM MgCl_2_, 0.1 mM EDTA] containing fatty-acid–free bovine serum albumin (BSA; 1 mg/mL) and ascorbic acid (0.1 mg/mL). Initial screening used a single concentration of 10 µM in in-plate quadruplicate. [^3^H]-LSD was used near its equilibrium dissociation constant (K_d_; final 0.5 nM). For each well, 25 µL compound (50 µM), 25 µL radioligand, and 75 µL membranes (5-HT_2A_ in binding buffer) were combined (final volume 125 µL). Nonspecific binding was defined in the presence of 10 µM clozapine. After a 2 h incubation at room temperature, receptor-bound radioactivity was collected by rapid filtration onto UniFilter-96 GF/C plates (PerkinElmer) using a FilterMate harvester; plates were dried, treated with MicroScint-O, and counted on a MicroBeta scintillation counter.

### Aminergic GPCR binding assay

Off-target binding across 21 aminergic GPCRs was performed at the NIMH PDSP using membranes from HEK293 cells transiently transfected with each receptor in a 96-well format, following target-specific PDSP protocols. Assays were run in triplicate with 12-point serial dilutions (0, 0.1, 0.3, 1, 3, 10, 30, 100, 300 nM; 1, 3, 10 µM) in a final volume of 125 µL/well of standard binding buffer [50 mM Tris-HCl (pH 7.4), 10 mM MgCl_2_, 0.1 mM EDTA]. Radioligand concentrations were selected close to target-specific K_d_ values. Total and nonspecific binding were defined by the absence or presence, respectively, of 10 µM reference ligand appropriate to each receptor. Following a 90 min incubation at room temperature, samples were processed identically to the 5-HT_2A_ assay. Equilibrium inhibition constants (K_i_) were calculated from fitted IC_50_ values using the Cheng–Prusoff equation, accounting with the assay-specific radioligand concentration and K_d_.

### Calcium mobilization assays

Intracellular Ca2+ flux was quantified with Fluo-4 Direct (Invitrogen) in HEK293 cell lines stably expressing 5-HT_2A_, 5-HT_2B_, or 5-HT_2C_. Cells were maintained in Dulbecco’s modified Eagle’s medium (DMEM) supplemented with 10% fetal bovine serum (FBS), penicillin (100 U/mL), streptomycin (100 µg/mL), hygromycin B (100 µg/mL), and blasticidin (15 µg/mL) at 37 C in 5% CO2. Receptor expression was induced with tetracycline (1.5 µg/mL), after which cells were seeded into poly-L-lysine–coated, black, clear-bottom 384-well plates at ∼15,000 cells/well in DMEM + 1% dialyzed FBS.

Twenty-four hours later, cells were loaded with 20 µL/well Fluo-4 Direct supplemented with 2 mM probenecid (Thermo Fisher Scientific) in drug buffer [HBSS, 20 mM HEPES, 0.1% (w/v) BSA, 0.01% (w/v) ascorbic acid, pH 7.4] for 1 h at 37 C/5% CO2, followed by 10 min at room temperature in the dark. Drug solutions were prepared at 3x in drug buffer and transferred to a 384-well source plate for additions yielding a 1x final concentration (3 µM). Fluorescence was recorded on a FLIPR-Tetra (Molecular Devices): a 10 s baseline (1 read/s) was followed by 120 s post-addition acquisition (1 read/s) after dispensing 10 µL of 3x drug. For antagonist mode, 10 µL of 5-HT (10 nM final) was added 15 min after test-compound addition. For analysis, the maximum fluorescence within 120 s was transformed to fold change over baseline (mean of the first 10 reads), normalized to 5-HT (agonist control) or clozapine (antagonist control), and plotted in GraphPad Prism (v10.1.0). An activity threshold of ≥10% of the maximal 5-HT response (agonism) or ≥20% of the inhibitory effect produced by clozapine (antagonism) was used to identify compounds that exhibit agonist or antagonist activity, respectively.

### PRESTO-Tango GPCR-ome screening

GPCR-wide selectivity of the selected compounds was profiled by measuring β-arrestin recruitment with PRESTO-Tango, using minor adaptations to established workflows (PMID: 25895059; 36173843; 37137306). HTLA cells were seeded at 10,000 cells/well in 40 µL DMEM + 1% dialyzed FBS in poly-L-lysine–coated, white, clear-bottom 384-well plates. After ∼6 h, cells were transfected with 20 ng plasmid DNA/well and incubated overnight. The next day, 10 µL of test compound (in DMEM + 1% dialyzed FBS) was added, and plates were incubated overnight. Media were then removed, and 20 µL/well of Bright-Glo reagent diluted in assay buffer [HBSS, 20 mM HEPES (pH 7.4)] was added. After a 20 min incubation at room temperature in the dark, luminescence was recorded.

Each plate included dopamine D_2_ receptor (DRD2) stimulated with 100 nM quinpirole as a positive control. Each GPCR was tested in quadruplicate at 0 and 10 µM (final), and responses were reported as fold change relative to the 0 µM condition.

## Associated Content

### Supporting information

Supporting information is available free of charge on the ACS Publication website.

**Table S1:** List of aminergic GPCRs and related targets used in SEA analysis

**Figure S1:** SEA predictions using a stricter p-value cutoff of 10^-25^ for significance

**Figure S2:** SEA predictions using a stricter p-value cutoff of 10^-50^ for significance

**Figure S3:** SEA predictions for all targets beyond aminergic set

**Figure S4:** Calcium mobilization hits with stricter definitions of agonism and antagonism

**Figure S5:** GPCR-ome hits with a stricter definition of off-target activity

**Data S1:** DOCK scores for top 10,000 molecules from in-stock and ZINC screens

**Data S2:** DOCK scores for molecules experimentally tested from in-stock and ZINC screens

**Data S3:** All SEA predictions for top 10,000 molecules from in-stock and ZINC screens

**Data S4:** All SEA predictions for experimentally tested molecules from in-stock and ZINC screens

**Data S5:** Primary binding for experimentally tested in-stock molecules against 5-HT_2A_

**Data S6:** Calcium mobilization for experimentally tested in-stock molecules against 5-HT_2_ receptor family

**Data S7:** GPCR-ome activity for experimentally tested in-stock and ZINC molecules

**Data S8:** Primary binding for experimentally tested in-stock and ZINC molecules against a subset of aminergic GPCRs and related targets

**Data S9:** pKis for experimentally tested in-stock and ZINC molecules against a subset of aminergic GPCRs and related targets

## Author Contributions

B.L.R conceived the project. J.J.I did in-stock database curation and preparation.

B.W.H did the in-stock docking, SEA calculations, data analysis, and figure making.

K.S. and X.P.H. designed and executed the GPCR-ome, radioligand-binding, and calcium-flux experiments, performed data analyses, and drafted the relevant sections of the manuscript. B.L.R and B.K.S supervised the project. B.W.H wrote the paper with input from all authors and primary editing by B.K.S and B.L.R.

## Acknowledgments

We thank Jiankun Lyu for helping locate and collect ultra-large 5-HT_2A_ docking results. Receptor binding profiles was generously provided by the National Institute of Mental Health’s Psychoactive Drug Screening Program, Contract # 75N95023C00021 (NIMH PDSP). The NIMH PDSP is Directed by Bryan L. Roth MD, PhD at the University of North Carolina at Chapel Hill and Project Officer Jamie Driscoll at NIMH, Bethesda MD, USA.

## Notes

The authors declare the following competing financial interests: B.K.S. is cofounder of BlueDolphin LLC, Epiodyne Inc, and Deep Apple Therapeutics, Inc., serves on the SRB of Genentech, the SABs of Schrödinger LLC and of Vilya Therapeutics, and consults for Frontier Discovery. J.J.I. is a cofounder of BlueDolphin LLC and of Deep Apple Therapeutics. B.L.R. is on the SABs of XyloBio, Epiodyne and Septerna and a cofounder of ImprintBio. The authors declare no other competing interests.

## Data and Software Availability

All docking results, docking poses, and experimental data are made freely available at lsd.docking.org. SEA calculations can be run at tldr.docking.org.

## Table of Contents Graphic

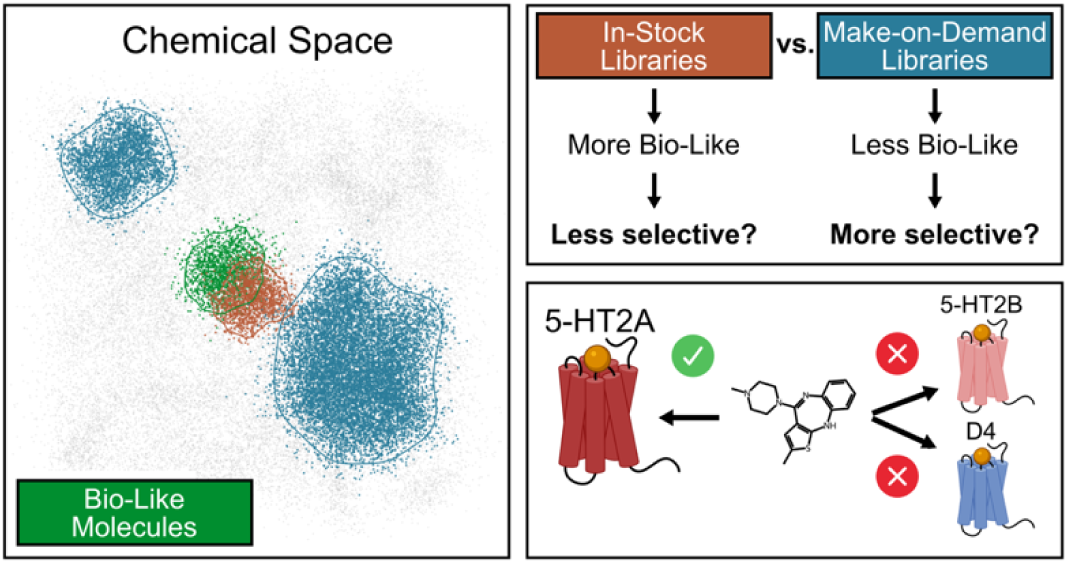

