## Supplemental Tables and Figures for "The selectivity implications of docking libraries with greater and lesser similarities to bio-like molecules"

**ASSOCIATED CONTENT**

**Target**

ACHE

ADRA1A

ADRA1B

ADRA1D

ADRA2A

ADRA2B

ADRA2C

ADRB1

ADRB2

ADRB3

CCR1

CCR2

CCR3

CCR4

CCR5

CCR6

CCR7

CCR8

CHRNA1

CHRNA2

CHRNA3

CHRNA4

CHRNA5

CHRNA6

CHRNA7

CHRNA8

CHRNA9

CHRNA10

CHRM1

CHRM2

CHRM3

CHRM4

CHRM5

CXCR1

CXCR2

CXCR3

CXCR4

CXCR5

CXCR6

CXCR7 (ACKR3)

DRD1

DRD2

DRD3

DRD4

DRD5

HRH1

HRH2

HRH3

HRH4

HTR1A

HTR1B

HTR1D

HTR1E

HTR1F

HTR2A

HTR2B

HTR2C

HTR3A

HTR3B

HTR3C

HTR3D

HTR3E

HTR4

HTR5A

HTR6

HTR7

OPRD1

OPRK1

OPRM1

SLC6A2

SLC6A3

SLC6A4

SIGMAR1

SIGMAR2

TAAR1

TAAR2

**Table S1.** List of aminergic GPCRs and other related proteins used for SEA analysis.

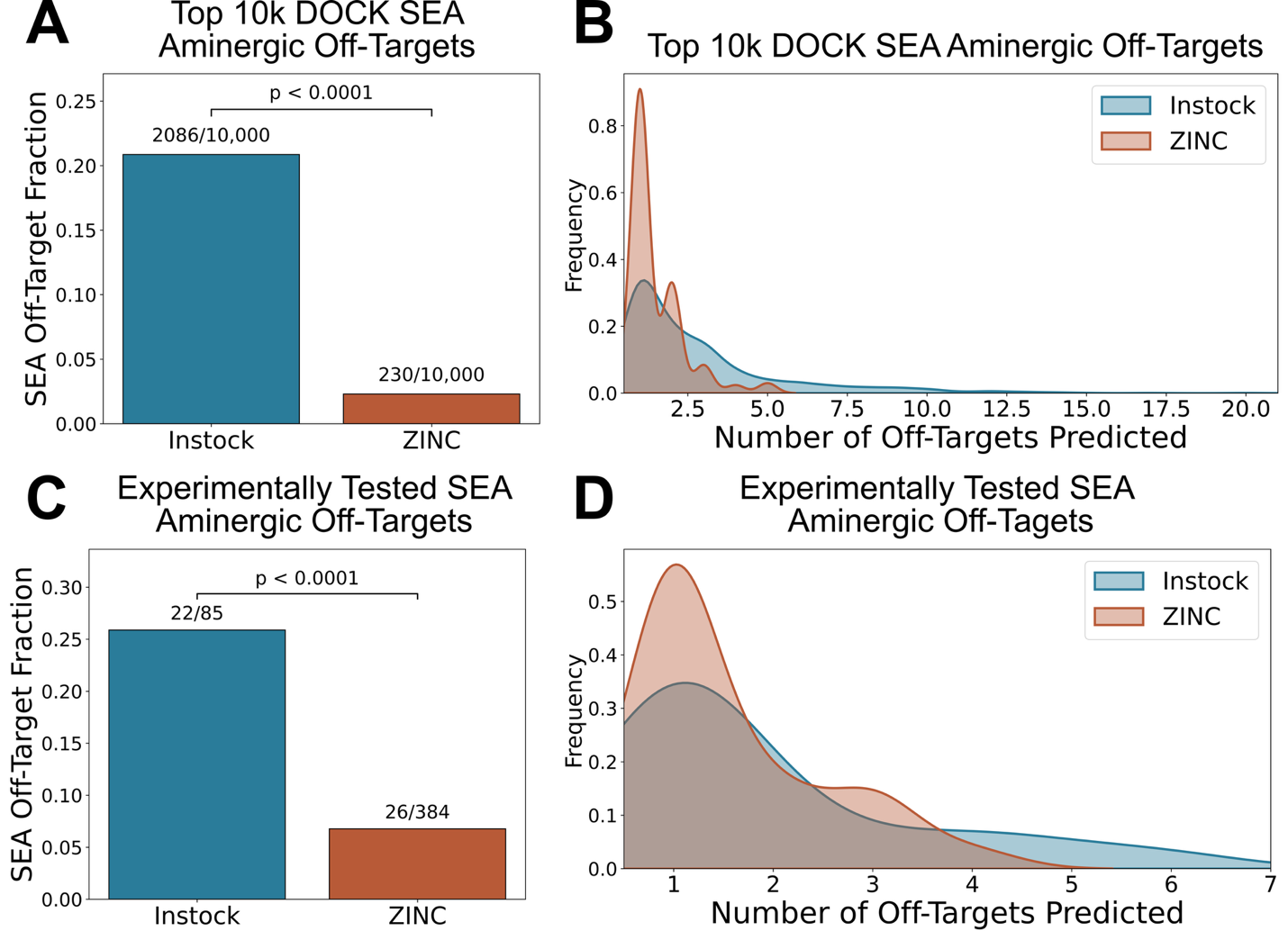

**Figure S1. SEA predictions with p-values less than 10^-25^. (A)** Fraction of the top 10,000 DOCK-scoring molecules predicted by SEA to bind any aminergic GPCR. **(B)** Number of SEA off-target predictions for the top 10,000 molecules. **(C)** Fraction of experimentally tested molecules predicted to bind any aminergic GPCR. **(D)** Number of SEA off-target predictions for the experimentally tested molecules.

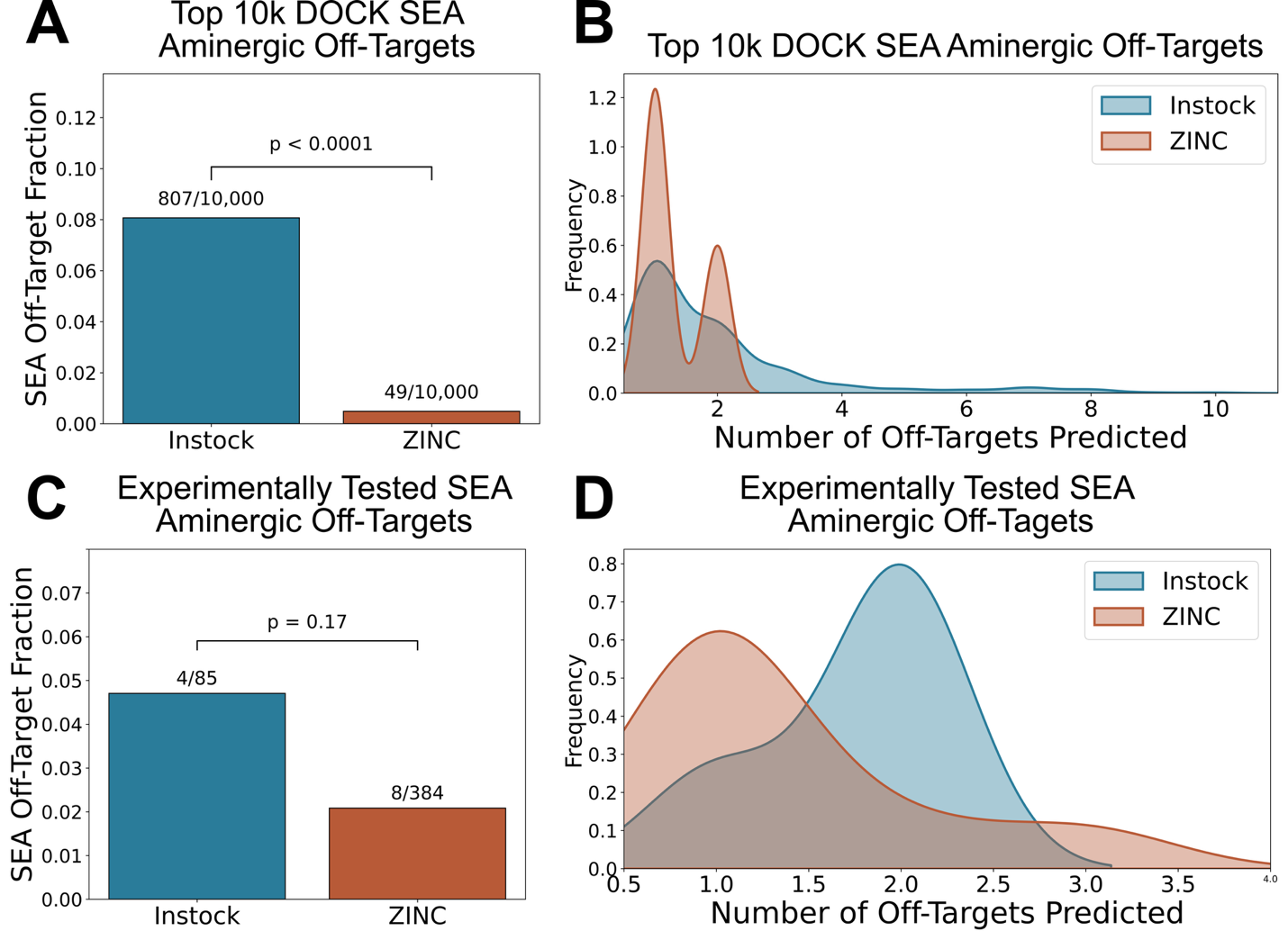

**Figure S2. SEA predictions with p-values less than 10^-50^. (A)** Fraction of the top 10,000 DOCK-scoring molecules predicted by SEA to bind any aminergic GPCR. **(B)** Number of SEA off-target predictions for the top 10,000 molecules. **(C)** Fraction of experimentally tested molecules predicted to bind any aminergic GPCR. **(D)** Number of SEA off-target predictions for the experimentally tested molecules.

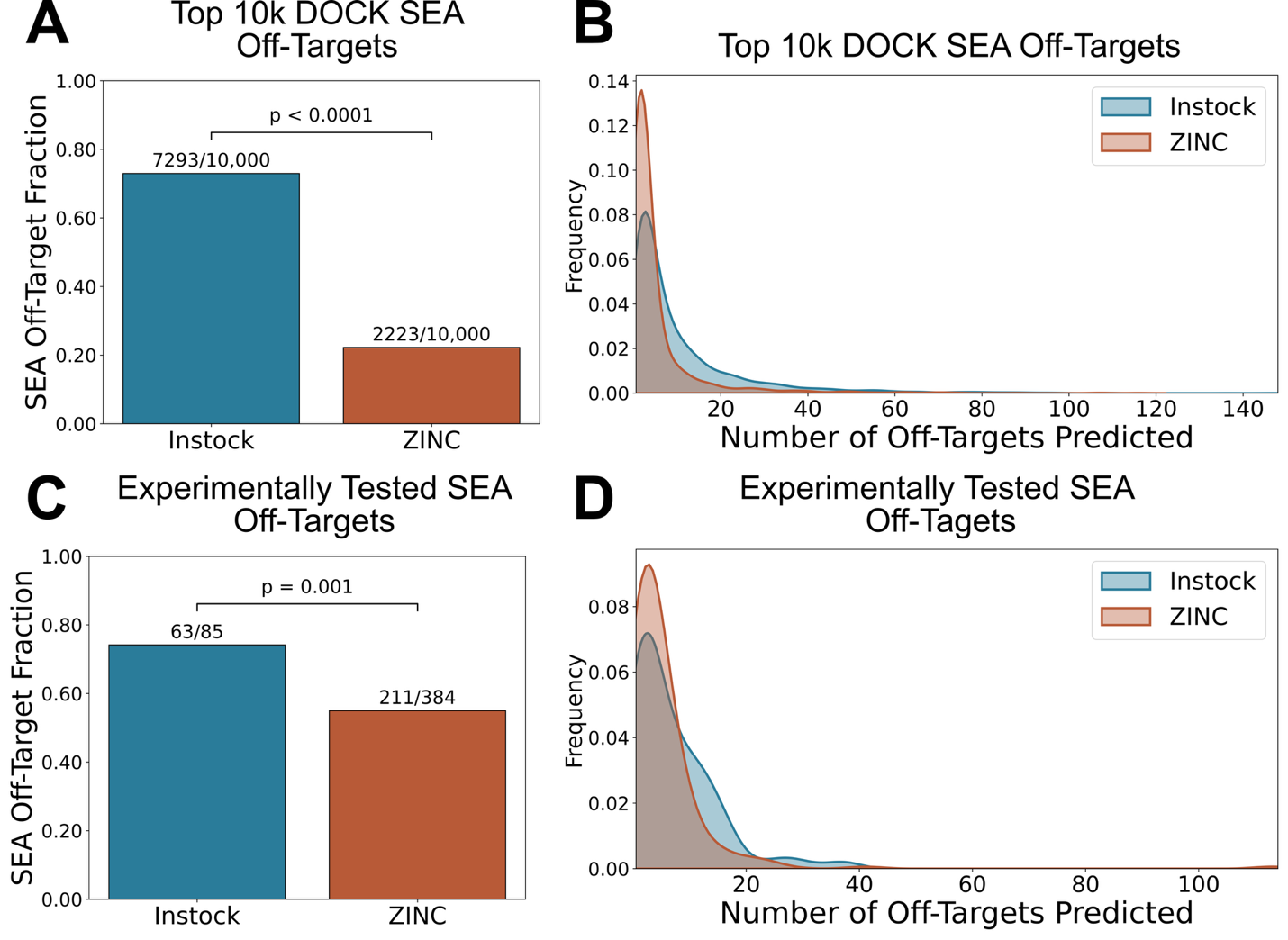

**Figure S3. SEA predictions with p-values less than 10^-10^ for all targets beyond aminergic GPCRs. (A)** Fraction of the top 10,000 DOCK-scoring molecules predicted by SEA to bind any target. **(B)** Number of SEA off-target predictions for the top 10,000 molecules. **(C)** Fraction of experimentally tested molecules predicted to bind any target. **(D)** Number of SEA off-target predictions for the experimentally tested molecules.

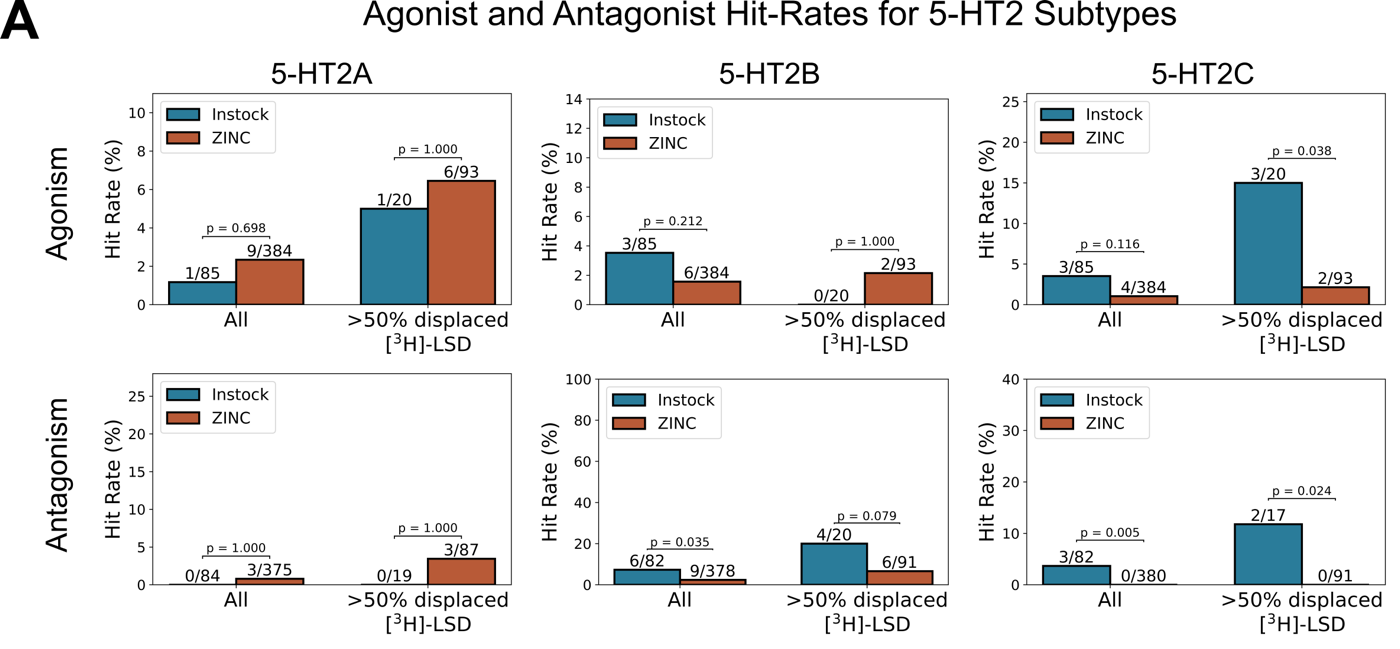

**Figure S4. 5-HT_2A/2B/2C_ calcium mobilization assays with stricter agonism and antagonism definitions. (A)** Agonist and antagonist hit rates for in-stock and ZINC molecules vs. 5-HT_2A_, 5-HT_2B_, 5-HT_2C_. Agonist hits were defined as >20% of 5-HT activity; antagonist hits were defined as >40% of clozapine activity.

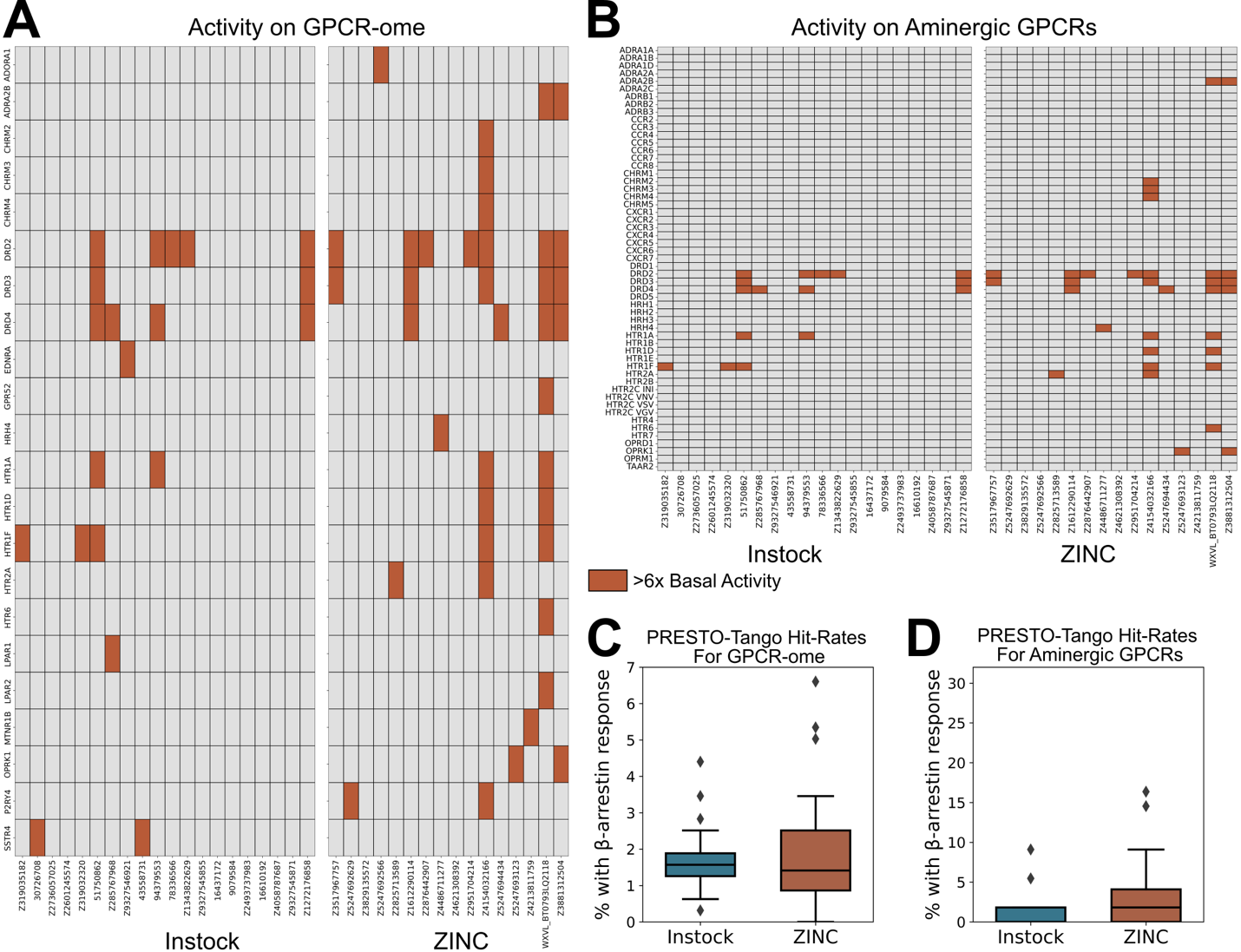

**Figure S5. Druggable GPCR-ome activity with stricter hit definition.** PRESTO-Tango hits for in-stock and ZINC molecules across the **(A)** druggable GPCR-ome and the **(B)** aminergic GPCRs. Molecules were considered hits if they produced a >6-fold increase in activity over basal levels. PRESTO-Tango hit rates for in-stock and ZINC molecules across the **(C)** druggable GPCR-ome and the **(D)** aminergic GPCRs.
